# Active and Passive Mechanical Deficits Precede Spinal Curvature in a Zebrafish Model of Idiopathic Scoliosis

**DOI:** 10.64898/2026.04.29.721663

**Authors:** Johnathan R. O’Hara-Smith, Samuel G. Bertrand, Julissa Ortiz-Delatorre, Rachael M. Giersch, Luke A. Rethwill, Damien M. Callahan, Daniel T. Grimes

**Author notes:** co-corresponding authors: DMC DTG.

## Abstract

Idiopathic scoliosis is a common spinal disorder characterized by progressive three-dimensional curvature of unknown cause. Although biomechanical imbalance has long been proposed to contribute to scoliosis, the early physiological states that precede curvature onset remain poorly understood. Here, we investigated this problem using zebrafish *uts2r3* mutants, which develop fully penetrant juvenile-onset spinal curvature following disruption of urotensin signaling. Transcriptomic analysis before curvature revealed altered expression of muscle-associated genes, suggesting that Uts2r3 influences axial muscle development or function. However, immunofluorescence, birefringence imaging, and quantitative analysis of myotome morphology showed that mutants lack overt muscle architectural defects or dystrophic pathology. By contrast, direct measurements of isolated larval trunks revealed pre-curvature biomechanical abnormalities: namely, *uts2r3* mutants generated reduced active force following electrical stimulation while also exhibiting increased passive resistance to stretch. These findings identify urotensin signaling as a regulator of axial tissue biomechanics during growth and suggest that scoliosis-like curvature can arise from an early imbalance between active force generation and passive tissue stiffness.

**Significance:** Spinal curvature is common, but the biological events that cause the spine to bend during growth remain poorly understood. Animal models, especially zebrafish, make it possible to study these events before curvature begins. Zebrafish lacking urotensin signaling develop spinal curves that arise during juvenile growth, similar to idiopathic scoliosis in humans. Here, we demonstrate that zebrafish lacking the urotensin pathway receptor Uts2r3 develop an abnormal biomechanical state prior to curve onset. Their axial tissues generate less active force when contracting and, at the same time, show increased passive resistance to stretch—an unexpected combination that reveals a distinct pre-curvature biomechanical state. These findings suggest that spinal curvature can arise from an early imbalance in tissue mechanics during growth and identify urotensin signaling as a pathway that helps preserve spinal morphology through a biomechanical mechanism.

## INTRODUCTION

The emergence and maintenance of body form is a fundamental problem in developmental biology. While early embryonic patterning establishes the basic organization of tissues, preserving correct geometry during growth requires continuous coordination between genetic programs and mechanical environments (1, 2). Failures in this coordination can produce progressive morphological defects with major functional consequences. A prominent example is idiopathic scoliosis (IS), a disorder characterized by rotational lateral spinal curvature that affects approximately 1-4% of populations and most often arises during adolescent growth (3). Despite its prevalence, the biological mechanisms that destabilize axial morphology over time remain poorly understood.

Spinal curvature has long been hypothesized to have a biomechanical origin arising from imbalances in forces acting on the growing spine (4). Classic models propose that axial alignment depends on a balance between active forces generated by musculature and passive resistance provided by connective tissues, including myofibrillar and tendinous elements and their connection to developing skeletal elements (5–7). Disturbances to this balance are thought to amplify asymmetries in loading and promote progressive curvature during growth (8). Consistent with this view, specific types of scoliosis are associated with neuromuscular or connective tissue disorders (9, 10) such as Duchenne muscular dystrophy (11). Current therapeutic strategies for IS, typically bracing and physiotherapeutic exercise, aim to redistribute mechanical loads (12, 13). However, direct tests of how defined genetic pathways linked to spinal curvature influence the mechanical properties of axial tissues during growth remain limited.

Zebrafish have emerged as a powerful model for investigating the origins of spinal curvature (1, 14). Key features of axial organization are conserved between zebrafish and humans, and many zebrafish mutants develop spinal curves that resemble IS (15–23). Previous work showed that disruption of motile cilia driven cerebrospinal fluid (CSF) flow within the brain ventricles and central canal leads to IS-like spinal curves during larval growth (23). Subsequent studies identified the urotensin II-related peptides, Urp1 and Urp2, as downstream effectors of CSF flow dependent control of body shape (21, 22, 24, 25). These peptides signal through the G protein-coupled receptor Uts2r3, and combined loss of *urp1* and *urp2*, or loss of *uts2r3*, causes juvenile onset spinal curvature in zebrafish (**Fig 1A**; (21, 22, 24)). These findings implicated urotensin signaling in the maintenance of spinal morphology during growth. Uts2r3 is expressed in skeletal muscle during zebrafish development (22), raising the hypothesis that this pathway influences spinal morphology by tuning trunk muscle biomechanics to coordinate the development of skeletal muscle with other rapidly growing tissues in embryonic and juvenile stages.

**Fig. 1.**
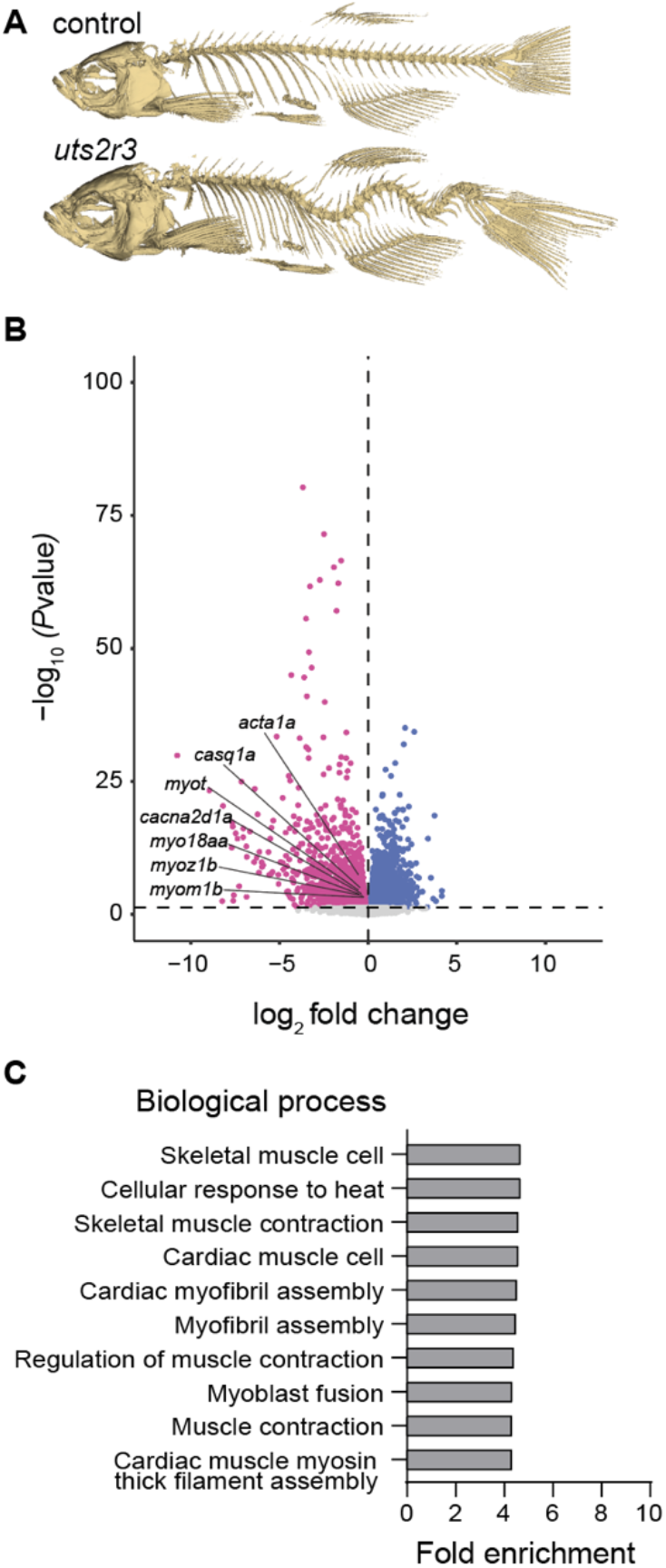
Loss of uts2r3 causes spinal curvature and alters muscle-associated gene expression before curve onset. *(A)* Representative skeletal preparations of adult sibling control and *uts2r3* mutant zebrafish. *uts2r3* mutants develop spinal curvature during juvenile growth. *(B)* Volcano plot of differential gene expression from RNA sequencing of 28 h.p.f. *uts2r3* mutants and wild-type siblings. Highlighted genes are associated with muscle development, myofibrillar organization, contraction, or calcium-dependent physiology. Dashed lines indicate significance and fold change thresholds. *(C)* Gene ontology biological process enrichment analysis of differentially expressed genes, showing enrichment of muscle-associated terms among genes downregulated in *uts2r3* mutants.

Here, we tested this hypothesis by combining transcriptomic analysis, imaging of muscle organization, and direct physiological measurements of active and passive mechanical properties in zebrafish *uts2r3* loss of function mutants. By assessing these features before the onset of spinal curvature, we sought to identify early biomechanical defects that could predispose the growing spine to develop the abnormal curvature typical of *uts2r3* mutants (**Fig. 1A**, (21, 24)). We found that loss of *uts2r3* alters muscle associated gene expression, reduces active force generation elicited from muscle, and increases passive elastic and viscous modulus, despite largely preserved muscle architecture. Together, these results identify urotensin signaling as a pathway that helps maintain spinal morphology by regulating the mechanical state of axial tissues during growth.

## RESULTS

### Muscle-related gene expression is altered in *uts2r3* mutants

During larval axial morphogenesis, motile cilia-dependent signaling promotes expression of *urp1* and *urp2* in the central canal (21, 22, 24). The Urp1/Urp2 receptor, Uts2r3, is expressed in skeletal muscle (22), suggesting that Urp1/Urp2 might signal directly to muscle tissue. We therefore hypothesized that Urp1/Urp2-Uts2r3 signaling contributes to the maintenance of axial morphology during juvenile growth by modulating muscle development or function (21, 24).

To test this hypothesis, we first performed bulk RNA-sequencing on 28 hours post fertilization (h.p.f.) *uts2r3* loss-of-function mutants and wild-type siblings, using five biological replicates per condition. We then identified genes that were significantly differentially expressed (*padj* < 0.05; **Fig. 1B**). Several muscle-associated genes were downregulated in *uts2r3* mutants, including myosin and skeletal muscle alpha-actin family members (*myo18aa, myo18ab, acta1a, acta1b*), genes associated with myofibrillar structure or myogenesis (*myom1b, myot, myoz1b*), and genes linked to calcium handling or excitable cell physiology (*casq1a, cacn1ha*). Gene ontology enrichment analysis supported this pattern, revealing overrepresentation of muscle-related terms including “skeletal muscle cell,” “myofibril assembly,” and “skeletal muscle contraction” among differentially expressed genes (**Fig. 1C**). These sequencing results indicate that loss of Uts2r3, a muscle-expressed Urotensin pathway receptor, leads to mis-regulation of muscle-associated genes prior to the onset of spinal curvature.

### *uts2r3* mutants lack overt structural defects but show subtle disruption of muscle organization

To assess whether transcriptional changes in *uts2r3* mutants were accompanied by altered muscle architecture, we examined trunk muscle using F-actin labeling and dystrophin immunofluorescence. At 24, 28, and 32 h.p.f., sarcomeric actin organization and dystrophin localization appeared normal in *uts2r3* mutants relative to wild-type siblings, with no obvious disruptions in fiber continuity or segmental patterning (**Fig. 2A** and **Fig. S1**). These results indicate that early transcriptional changes observed in *uts2r3* mutants are not accompanied by gross defects in larval muscle architecture.

**Fig. 2.**
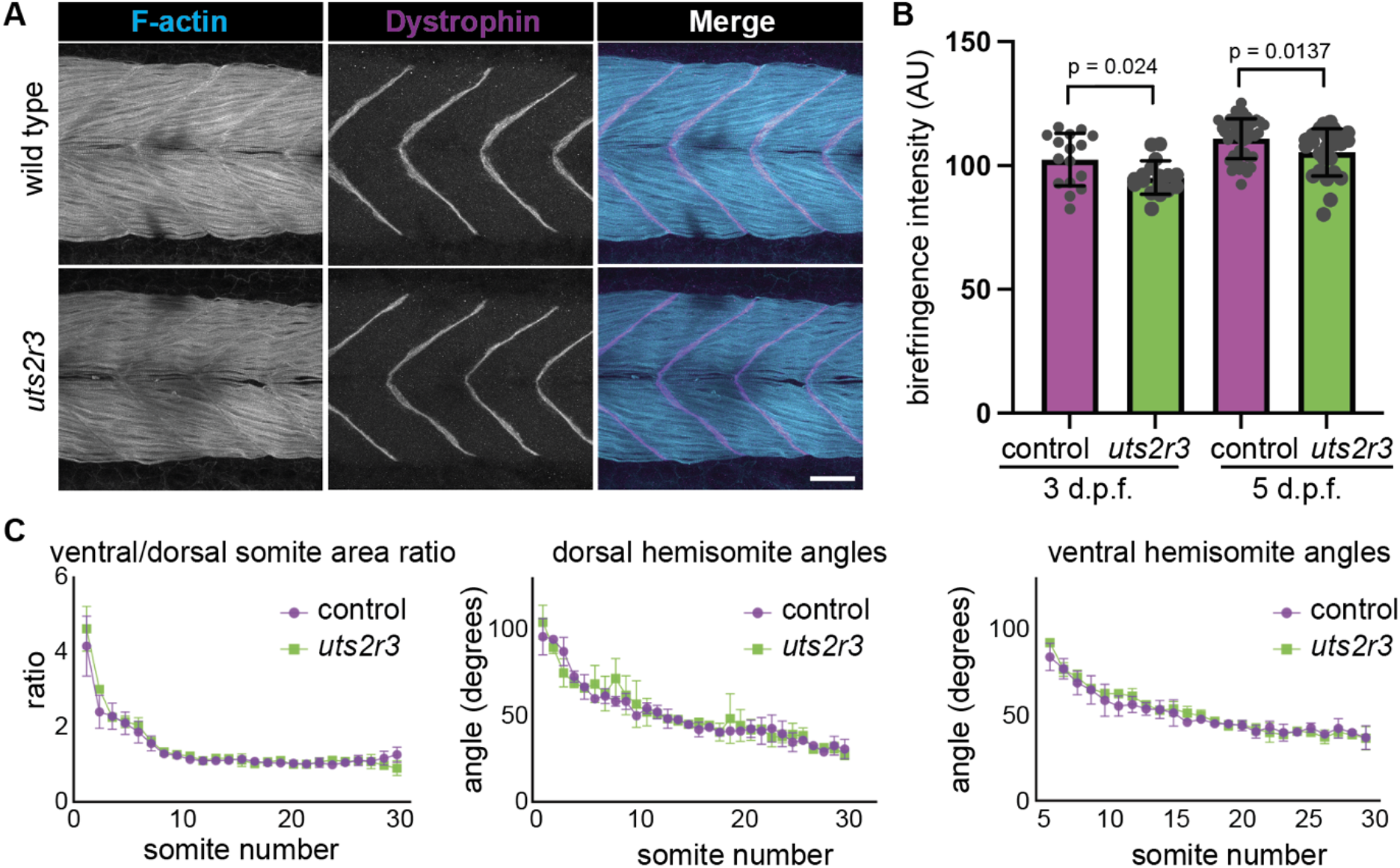
*uts2r3* mutants show subtle disruption of muscle organization without overt dystrophic pathology or dorsoventral hemisomite asymmetry. *(A)* Representative trunk muscle images from 32 h.p.f. wild-type and *uts2r3* mutant embryos labeled for F-actin and dystrophin. F-actin organization and dystrophin localization appear grossly preserved in *uts2r3* mutants, with no obvious disruption of muscle fiber continuity or myoseptal patterning. Images show equivalent trunk regions acquired at 40x magnification. Scale bar, 20 μm. N ≥ 3 fish examined per condition. *(B)* Quantification of trunk muscle birefringence intensity in 3 d.p.f. and 5 d.p.f. larvae. *uts2r3* mutants showed a modest but significant reduction in mean birefringence intensity at both stages. Data are mean ± SD with individual larvae shown as points. 3 d.p.f.: control, n = 16, *uts2r3*, n = 17. 5 d.p.f.: control, n = 44, *uts2r3*, n = 25. Statistical significance was determined by Student’s *t* test. *(C)* Quantification of dorsoventral hemisomite proportions and angles at 14 d.p.f. in sibling-matched control and *uts2r3* mutant larvae carrying the skeletal muscle reporter *Tg(unc503:mVenus-CAAX)*. Ventral-to-dorsal hemisomite area ratios were measured somite-by-somite along the trunk and tail. Dorsal and ventral hemisomite angles were measured relative to the midline. No significant differences were detected between genotypes. Data are mean ± SD. Control, n = 4 larvae; *uts2r3*, n = 3 larvae.

We next asked whether more subtle abnormalities emerge later during larval growth by quantifying trunk muscle birefringence. Because birefringence depends on the ordered alignment of myofibrils, reductions in signal provide a sensitive readout of myofibrillar disorganization or misalignment. At 3 and 5 days p.f. (d.p.f.), *uts2r3* mutants showed a modest but consistent reduction in birefringence intensity compared with wild-type siblings (**Fig. 2B**). However, birefringence patterns remained smooth and continuous, and we did not observe breaks or gaps characteristic of dystrophic muscle mutants (26, 27**; Fig. S2**). These data indicate mild disruption of muscle organization without overt dystrophic pathology.

Because asymmetric muscle size and mass have been studied in human contexts as a potential feature underlying or perhaps even driving IS, we next tested whether dorsoventral muscle asymmetry was altered in *uts2r3* mutants (**Fig. S3**). To label myotome morphology, we generated Tg(*unc503:mVenus-CAAX*) transgenic fish in which the muscle-specific *unc45b* promoter drives mVenus expression in skeletal muscle. Using this line, we measured dorsal and ventral hemisomite area at 14 d.p.f., just as ventrally-directed spinal curves became apparent (24). Despite axial curvature, the ratio of ventral to dorsal hemisomite area in *uts2r3* mutants was indistinguishable from that of sibling controls (**Fig. 2C**). Hemisomite angles relative to the midline were also unchanged, suggesting that myosepta orientation is preserved in mutants (**Fig. 2C**). These results argue against overt dorsoventral asymmetry in muscle growth as the cause of spinal curvature.

Taken together, these data show that loss of Uts2r3 causes early transcriptional changes and later subtle disruption of muscle organization without gross defects in muscle architecture, dystrophic pathology, or asymmetric muscle growth.

### Active muscle force is reduced in *uts2r3* mutants

The subtle structural abnormalities observed in *uts2r3* mutants raised the possibility that urotensin signaling influences inherent muscle function rather than gross muscle architecture or morphology. We therefore asked whether loss of *uts2r3* impairs the ability of axial muscle to generate active force. To test this directly, we measured force production in isolated larval trunk preparations using an apparatus for microscale measurements of force and displacement in contractile tissues that was adapted for use in zebrafish larvae (**Fig. 3A**). Trunk segments were mounted between a force transducer and a length motor using paired sutures (**Fig. 3A**) attaching the specimens to stainless steel hooks, allowing precise measurements of electrically evoked contraction while preserving native tissue organization. Because muscle activation was achieved via an external electrical stimulus, this preparation isolates intrinsic mechanical properties of the muscle while minimizing contributions from motor neurons or locomotor behavior.

**Fig. 3.**
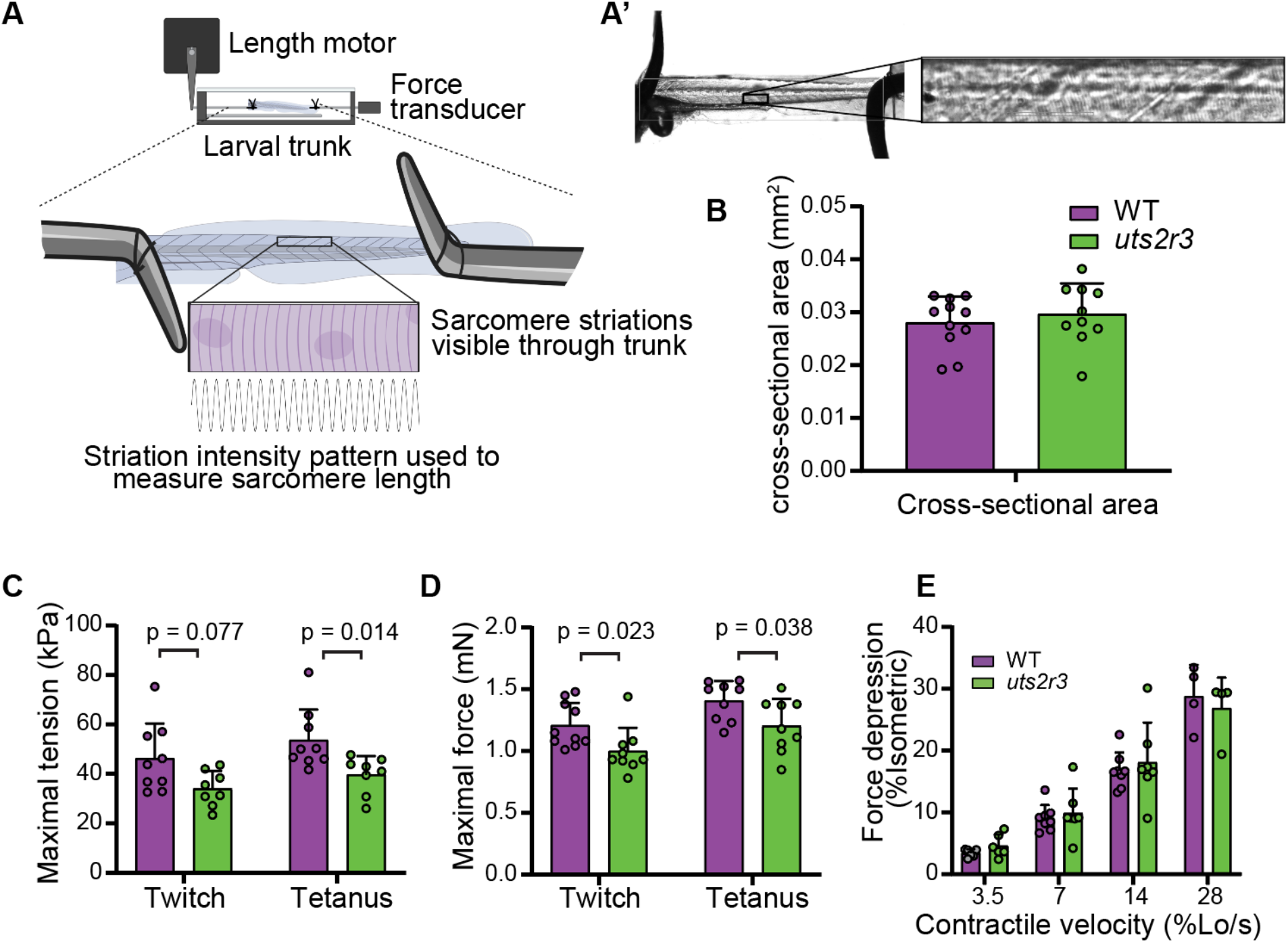
Active force generation is reduced in 5 d.p.f. *uts2r3* mutant larval trunks. *(A)* Schematic of the *ex vivo* larval trunk mechanics assay. Isolated 5 d.p.f. preparations were mounted between a length motor and force transducer for mechanical testing. Sarcomere striations were visualized and used to measure muscle length before mechanical assessment. *(A’)* Representative image of a mounted 5 d.p.f. trunk preparation, with the expanded view showing sarcomere striations. *(B)* Cross-sectional area of larval trunk preparations, calculated from dorsal-ventral and lateral diameter measurements assuming an elliptical cross section. Cross-sectional area did not differ between wild-type and *uts2r3* mutant specimens (*p* = 0.495; WT, n = 10; *uts2r3*, n = 11). *(C)* Maximal tension generated during twitch and tetanic stimulation. *uts2r3* mutants generated reduced tension compared with wild type, with a significant reduction during tetanic stimulation. WT, n = 9; *uts2r3*, n = 10. *(D)* Maximal force generated during twitch and tetanic stimulation. *uts2r3* mutants generated significantly less force under both stimulation conditions. WT, n = 9; *uts2r3*, n = 10. *(E)* Force depression during shortening contractions, expressed as percent reduction from isometric force at the same length. No significant differences were detected across contractile velocities. WT, n = 8; *uts2r3*, n = 7. Bars show means ± SD with individual specimens shown as points.

Larvae were analyzed at 5 d.p.f., before the onset of spinal curvature but after the emergence of transcriptional changes and reduced muscle birefringence (**Fig. 1B** and **2B**). To determine whether potential differences in force production could be explained by body size or gross morphology, we measured axial dimensions at three standardized positions along the trunk and tail. Axial dimensions did not differ significantly between *uts2r3* mutants and wild-type siblings (*p* = 0.495; **Fig 3B**). All force measurements (mN) were nevertheless normalized to average specimen cross sectional area to report tension normalized to size (mN·mm^-2^).

Using this preparation, we measured active force generation under two stimulation paradigms. Single twitch contractions, elicited by a brief (400 µs) electrical pulse, report force generated during an individual contraction cycle, whereas tetanic stimulation at high frequency (300 Hz) produces sustained contraction that elicits the maximal force-generating capacity of the muscle. Under both conditions, *uts2r3* mutants generated less force compared with wild-type siblings (**Fig. 3C and 3D**). This reduction persisted after normalization to axial dimensions (tetanic tension, *p* = 0.008; **Fig. 3C**), indicating that the deficit reflects reduced intrinsic force-generating capacity rather than differences in specimen size. The reduction was greatest under tetanic conditions, with tetanic tension decreased by approximately 35% in *uts2r3* mutants relative to wild type (*p* < 0.05; **Fig. 3C**).

The reduction in active force observed in *uts2r3* mutants could arise from several distinct mechanisms. Muscle weakness can reflect fewer or weaker force-producing crossbridges (binding events between filamentous actin and the molecular motor, myosin), but it can also result from altered contractile dynamics, such as slowed activation, impaired force transmission across parallel and serial elastic elements (i.e., connective tissue), or exaggerated force loss during shortening. To distinguish between these possibilities, we examined multiple parameters describing the kinetics of muscle contraction using the same intact larval preparation. First, we assessed the relationship between submaximal and maximal force production by quantifying the ratio of twitch force to tetanic force. A twitch contraction elicited by a single 400 µs stimulus reflects force-generating capacity of the tissue and rapid response of its intracellular components to activation and effective transmission of that force through non-contractile tissues. By contrast, tetanic stimulation produces sustained maximal activation and is therefore less reliant on dynamic processes that remain in flux throughout the activation period. Comparing twitch and tetanic force therefore helps reveal whether slowed myosin dynamics or impaired force transmission, which would disproportionately reduce twitch compared to tetanic force, are playing a role in our outcomes. This ratio did not differ between wild-type (0.87 ± 0.11) and *uts2r3* mutant (0.82 ± 0.13; *p* = 0.389), indicating preserved scaling between submaximal and maximal activation. Thus, although overall force output is reduced in *uts2r3* mutants, the efficiency with which force is transferred during activation is normal.

Second, the half-relaxation time of twitch contractions, defined as the time required for force to decay from its peak to 50% of peak, was indistinguishable between wild-type (10.3 ± 0.9 ms) and *uts2r3* mutants (10.6 ± 2.4 ms; *p* = 0.719). This parameter is sensitive to the rates of calcium reuptake and crossbridge detachment and therefore reflects the temporal control of contraction and relaxation. This result therefore indicates normal relaxation kinetics in *uts2r3* mutants.

Finally, because muscle normally shortens while generating force during locomotion, we asked whether *uts2r3* mutant muscle exhibits abnormal force loss during active shortening. In skeletal muscle, force output normally decreases as shortening speed increases, reflecting limits on how rapidly myosin crossbridges can generate force. We quantified this relationship by measuring force-velocity curves during isovelocity contractions across a range of shortening speeds. Across all velocities tested, shortening-dependent force loss was not different between wild type and *uts2r3* mutants (multivariate ANOVA, *p* > 0.1 for all comparisons; **Fig. 3E**), indicating preserved shortening-dependent modulation of force. Together, these data show that *uts2r3* mutants exhibit a pre-curvature deficit in active force generation despite preserved contractile kinetics.

### Passive stiffness of axial tissues is increased in *uts2r3* mutants

Active muscle force represents only one component of the mechanical environment experienced by the developing spine. Forces generated by muscle are transmitted through surrounding tissues, including muscle connective tissue, myosepta, and extracellular matrix, whose passive mechanical properties influence how those forces are distributed, resisted, and dissipated. In addition, muscle tissue itself has intrinsic viscoelastic properties that contribute to trunk mechanics when it is not actively contracting. We therefore asked whether loss of *uts2r3* alters the passive viscoelastic properties of axial tissues independent of active contraction.

To test this, we measured passive mechanical behavior in intact 5 d.p.f. larval trunks using a stretch-and-hold assay performed in the absence of electrical stimulation on the same force-length apparatus used for active force measurements. By adjusting the lengthener, preparations were subjected to repeated increments of axial strain, each corresponding to 2% of the initial specimen length, followed by a 2 min hold period to allow stress relaxation before the next stretch was applied. This sequence was repeated until specimens reached 156% of their initial length. This approach allowed us to distinguish elastic modulus (“stiffness”), reflected in the stress measured following stress-relaxation, from viscous modulus (“viscosity”), reflected in the time-dependent decay in stress during the hold phase (**Fig. 4A**).

**Fig. 4.**
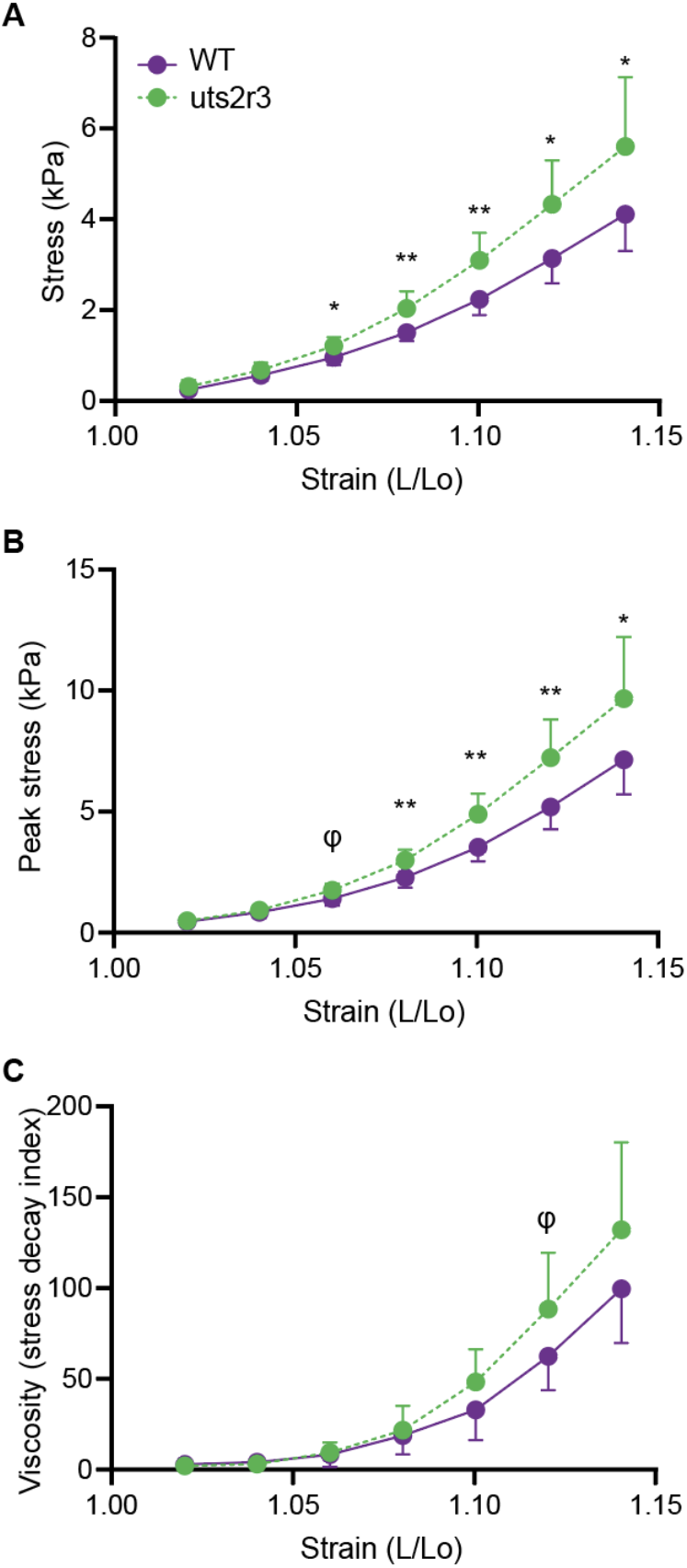
Passive mechanical resistance is increased in *uts2r3* mutant axial tissues. *(A)* Stress-relaxed passive stress measured during stretch-and-hold testing of isolated 5 d.p.f. larval trunk preparations. *uts2r3* mutants showed increased passive stress compared with wild-type siblings, particularly at higher strain levels. *(B)* Peak stress measured immediately after each stretch was also elevated in *uts2r3* mutants. *(C)* Viscous behavior, quantified using the stress decay index during the hold phase, showed a modest trend toward increased viscosity in *uts2r3* mutants. Data show group means ± SD. WT, n = 8; *uts2r3*, n = 5. φ, p < 0.1; *, p < 0.05; **, *p* < 0.01.

Using this assay, we found that axial tissues from *uts2r3* mutants exhibited increased passive stiffness compared with wild-type siblings. At low strain levels, mechanical behavior was similar between genotypes. At higher strains, however, *uts2r3* mutants displayed significantly elevated passive stress, corresponding to an approximately 35 to 40% increase in modulus at the highest strain tested (multivariate ANOVA at individual strain levels, *p* < 0.05; **Fig. 4A**). This difference was evident both in peak response measured immediately after stretch (**Fig. 4B)** and in stress values measured after relaxation following the hold phase, indicating a robust increase in elastic resistance to deformation in mutant tissues.

Passive viscous behavior, assessed by the time constant of stress relaxation during the hold phase, exhibited greater variability across samples (**Fig. 4C)**. Nevertheless, a consistent trend toward increased resistance to prolonged deformation was observed in *uts2r3* mutants (**Fig. 4A-C**). These results show that loss of Uts2r3 increases passive mechanical resistance of axial tissues before spinal curvature begins. Together, our findings show that Uts2r3 helps establish the balance of active and passive mechanical properties that maintain spinal morphology during growth.

## DISCUSSION

Our findings establish *uts2r3* mutants as a model in which altered biomechanics precede and are likely to contribute to IS-like spinal curvature. Before the onset of visible curvature, mutant larvae exhibit a striking combination of reduced active force generation and increased passive resistance to deformation, despite the absence of overt defects in muscle architecture or asymmetric muscle growth. These findings identify a previously unrecognized role for urotensin signaling in regulating the mechanical state of axial tissues during growth.

This mechanical phenotype provides a plausible context for the later emergence of spinal curvature in *uts2r3* mutants. Although it is still unclear whether reduced force generation or increased passive stiffness is sufficient to cause curvature, our data show that both abnormalities are present before curvature begins. We therefore favor a model in which loss of *uts2r3* destabilizes axial mechanics during growth, increasing susceptibility to pathological bending of the body axis. More broadly, these findings support the idea that spinal morphology depends on the continued maintenance of a balanced mechanical environment as the body grows.

*uts2r3* mutant axial muscle generates less active force despite largely preserved contractile dynamics. Twitch:tetanus relationships, relaxation kinetics, and force-velocity behavior were all broadly similar between mutants and wild-type siblings, arguing that the reduction in force is not accompanied by gross disruption of contraction timing or shortening-dependent force production. Instead, loss of *uts2r3* appears to reduce overall force-generating capacity while leaving several core physiological properties intact. This interpretation is consistent with our structural data, which show only subtle disruption of muscle organization rather than overt dystrophic pathology.

Loss of *uts2r3* also increases passive mechanical in axial tissues. In the absence of electrical stimulation, mutant trunks resisted stretch more strongly than those of wild-type siblings, especially at higher strain. Because passive mechanics in this preparation likely reflect contributions from muscle together with surrounding connective tissues, myosepta, and extracellular matrix, our data point towards a structural basis of this stiffening phenotype, although the specific tissues or combinations of tissues responsible remain to be identified. Thus, loss of *uts2r3* alters not only active force production but also the passive viscoelastic properties of the trunk.

Importantly, these active and passive defects are unlikely to be simple consequences of one another. Increased passive stiffness does not readily explain the reduction in active force generation, and the preservation of several contraction kinetics parameters argues against a generalized collapse of muscle function. Instead, *uts2r3* mutants exhibit a distinct mechanical imbalance in which the trunk is both weaker during active contraction and more resistant to passive deformation. Such a state would be expected to alter how forces are generated, transmitted, and absorbed along the growing body axis. We propose that this early imbalance in axial tissue mechanics contributes, directly or indirectly, to the later onset of IS-like curvature in *uts2r3* mutants.

Our results also complement recent work implicating altered tissue mechanics as an early feature of spinal curve pathogenesis in zebrafish. Pumputis et al. showed that oxidative stress induces intervertebral extracellular matrix remodeling, elevated spine stiffness, and later scoliosis-like curvature (28). Importantly, and as for our work, increased stiffness was detectable prior to curve onset (28). Together, these findings support the broader idea that spinal curvature can emerge from early disruption of the mechanical environment of the growing axis across distinct zebrafish models.

A key question raised by this work is how urotensin signaling regulates axial tissue mechanics at the cellular and molecular level. Uts2r3 is expressed in skeletal muscle, but the passive stiffening phenotype could reflect changes in muscle, extracellular matrix, myosepta, or interactions among these tissues. Future work defining the tissue-specific requirements for Uts2r3 and the downstream effectors that alter active and passive mechanics will be important for linking urotensin signaling to the maintenance of axial morphology.

More broadly, our findings argue that spinal curvature cannot be understood from genetic or cellular mechanisms alone. Instead, the effects of developmental pathways must ultimately be interpreted through the tissue-scale mechanical environments they create. Progress in this area will require closer interaction between genetic, cell biological, and biomechanical approaches, allowing us to understand how molecular changes are transformed into altered forces, stiffness, and progressive changes in body form during growth.

## Acknowledgements

We thank Judy Peirce, Tim Mason, and the Aquatics Facility for zebrafish husbandry, Adam Fries and the GC3F Biological Imaging Facility, and Autumn Wicklund and the Media Kitchen, all at the University of Oregon. This study was supported by National Institutes of Health grants R35GM142949 (to D.T.G.), R21HD117423 (to D.T.G.) and R21AG077125 (to D.M.C.).

## Author Contributions

J.O.S.: Conceptualization, Investigation, Data curation, Formal analysis, Visualization, Writing — original draft, Writing — review & editing. S.G.B.: Investigation, Data Curation, Formal analysis, Writing — review & editing. J.O.D.: Investigation, Data curation. R.M.G.: Investigation, Data curation. L.A.R.: Data curation. D.M.C.: Conceptualization, Funding acquisition, Investigation, Data curation, Formal analysis, Supervision, Visualization, Writing — original draft, Writing — review & editing. D.T.G.: Conceptualization, Funding acquisition, Supervision, Visualization, Writing — original draft, Writing — review & editing.

## METHODS

### Zebrafish

AB strain *D. rerio* were used for all experiments. The transgenic line generated in this study was *Tg*(*unc503:mVenus-CAAX)*^*b1513*^. We also used the previously published *uts2r3*^*b1436*^ line (24). All experiments were undertaken in accordance with research guidelines of the Association for Assessment and Accreditation of Laboratory Animal Care International and approved by the University of Oregon Institutional Animal Care and Use Committee.

### Birefringence quantification and analysis

Zebrafish larvae were placed directly on a polarizing light filter (K&F Concept, B0C2GWS95B) beneath a second polarizing filter (Educational Innovations, TRTV2091) mounted to a custom enclosure. Larvae were positioned with a deer hair (Ted Pella, Inc, 119) to maximize and standardize the birefringence signal. Images were acquired using identical settings on a Leica THUNDER stereoscope. Birefringence was quantified using FIJI (29) by measuring the mean pixel intensity across the contiguous birefringent region of the trunk.

### *Tg(unc503:mVenus-CAAX)* generation

The *Tg(unc503:mVenus-CAAX)* plasmid was generated using Multisite Gateway cloning. Entry clones p5E:*unc503*(Addgene, 64020), pME:*mVenus-CAAX* (Addgene, 75153), and p3E:polyA were combined with the Tol2 destination vector pDest:Tol2-polyA-cryaa:GFP (Addgene, 64022) using LR Clonase II (Ambion, 11791100). The assembly reaction was transformed into NEB 10-beta competent cells. Individual colonies were selected, plasmid DNA was isolated using the GeneJET Plasmid Miniprep Kit, and the full plasmid was sequenced by Plasmidsaurus to verify the final construct. Transgenesis injections were performed using 25 ng/µL plasmid DNA and 25 ng/µL Tol2 transposase mRNA. Tol2 mRNA was synthesized using the mMESSAGE mMACHINE kit (Thermo Fisher, AM1344).

### *Tg(unc503:mVenus-CAAX)* imaging

Stable single-copy integrations of Tg(*unc503:mVenus-CAAX*) were generated and used to visualize skeletal muscle morphology. Larvae carrying the transgene were mounted in 1% low-melting-point agarose (Fisher, BP165-25) in petri dishes containing coverslips (Mattek P35G-1.5-10-C). Images were acquired on a Zeiss 880 confocal microscope using a Plan-Apochromat 20x objective. Tiled images were collected with at least 10% overlap between adjacent tiles. Acquisition settings were configured in Zen imaging software using the eYFP channel to detect mVenus-CAAX fluorescence. Image tiles were converted to IMS format using Imaris File Converter 10.2.0 and stitched using Imaris Stitcher 9.9.0 to generate complete trunk images. Stitched images were converted to maximum intensity projections in FIJI. To quantify somite morphology, dorsal and ventral hemisomite boundaries were manually traced as regions of interest, and ROI areas were measured in FIJI. Ventral-to-dorsal hemisomite area ratios were then calculated for each somite. Myoseptal angles were measured from the midline along the visible trajectory of each myoseptum using the angle tool in FIJI.

### *uts2r3* genotyping by PCR

Genomic DNA was extracted from whole larvae, 14 d.p.f. fin clips, or adult fin tissue by alkaline lysis. Tissue was incubated in 50 mM NaOH (VWR, MK768004) for 20 minutes at 95 °C using 50 µL to digest whole larvae or 14 d.p.f. fin clips, and 100 µL for adult fin tissue. Samples were neutralized by adding 5 µL or 10 µL of 1M Tris, pH 7.5 (VWR 97061-794/VWR97061-258) to the 50 µL or 100 µL NaOH lysates, respectively. PCR genotyping was performed using three primers listed in the Key Resources Table and Taq DNA polymerase with standard buffer (NEB, M0273) for 44 amplification cycles. PCR products were resolved by electrophoresis on 1.5 to 2.0% agarose gels (Topvision Agarose, Thermofisher R0492) with DNA ladder. Genotypes were assigned based on allele-specific banding patterns, using known heterozygous control DNA to confirm successful amplification and band separation.

### Bulk RNA-sequencing and differential gene expression analysis

A DirectZol RNA kit (Zymo Research, R2050) was used to extract total RNA from 28 h.p.f. embryos. A Watchmaker RNA library Prep Kit (Watchmaker Genomics, 7K0078-024) was used for library preparation. Sequencing was performed on an Illumina NextSeq 2000, and trimming was performed using Cutadapt in R (30). Reads were aligned to a zebrafish transcript annotation (31) using the Rsubread package (32) and the resulting alignments were converted to BAM files. Further quality control and short-read quantification was conducted using QuasR (33). Counting of reads and differential gene expression analysis was performed using the DESeq2 package (34). A cutoff of < 0.05 was used for identifying significant hits based on adjusted p-value (*padj*). Gene ontology (GO) enrichment analysis was performed using ShinyGO 0.82 (35), incorporating the “Phenotype.GeneRIF.Predicted” pathway database to identify enriched biological processes among differentially expressed genes.

### Phalloidin/MANDRA1/DAPI Staining and Imaging

Embryos from 18-32 h.p.f. were fixed in 4% paraformaldehyde (PFA; Fisher Scientific, 50-980-492), then washed, permeabilized, and blocked. Embryos were incubated with a primary antibody solution containing MANDRA-1 supernatant (DSHB, MANDRA-1S) at a 1:50 dilution, followed by incubation with Hoechst 33258 nuclear stain (1:1000; Sigma-Aldrich, 94403), Phalloidin-AlexaFluor-488 (1:100, Invitrogen/Life Technologies, A12379), and donkey anti-mouse AlexaFluor-647 (1:250; ThermoFisher, A-31571). Embryos were mounted and imaged on a Zeiss LSM 880 confocal microscope.

### Active and Passive Mechanical Comparisons

Active contractile function and passive mechanical function were performed as previously described (36). Zebrafish larvae were reared at 28.5°C and maintained at this temperature until immediately before euthanasia. At 5 d.p.f., larvae were anesthetized, decapitated, and the swim bladders were removed using fine forceps (Fine Science Tools, 11255-20). Specimens were immediately transferred to oxygenated Ringer’s solution containing 2.5 mm CaCl_2_ (Sigma-Aldrich, 21115), 1.2 mM MgCl_2_ (Sigma-Aldrich, 208337), 1.2 mM KH_2_PO_4_ (ThermoFisher, P285), 11.1 mM glucose (TCI, G0048), 117.2 mM NaCl (Sigma-Aldrich, S9888), 4.7 mM KCl (Sigma-Aldrich, P9333), and 25.2 mM NaHCO_3_ (ThermoFisher, BP328-500). The solution was equilibrated to pH 7.4 with a mixture of 95% O_2_ and 5% CO_2_ and maintained at 18°C for subsequent assays.

Specimens were mounted to a micromanipulator and force transducer, typically used to measure single muscle fiber and muscle tissue composite specimens (Aurora Scientific, Aurora, ON, Canada). This apparatus was customized for these experiments with two stainless steel hooks that were used to mount each specimen at the caudal and rostral ends, secured with four sutures (**Fig. 3A**). Because zebrafish larvae are semi-translucent at this stage, sarcomere length was measured using an inverted microscope beneath the force-length apparatus. Once secured, specimens were adjusted to a sarcomere length of 1.9 µm, confirmed by observation of thick and thin filament striations.

Active contractile function was measured through field electrical stimulation by assessing the twitch-to-tetanus ratio and force-velocity relationships, as described previously (36). Twitch contractions were elicited using a single 400 µs stimulus, and tetanic contractions were elicited at 300 Hz. Force-velocity relationships were assessed during tetanic stimulation, with tension measured at velocities between 0 and 28 Lo/s. Stimulation was provided using a constant-current stimulation device (Aurora Scientific), with supramaximal current determined independently for each sample preparation before mechanical assessment.

Passive viscoelastic properties, including viscous and elastic modulus, were assessed using stretch-and-hold assays. For these assays, the starting length was adjusted to a sarcomere length of = 1.8 µm. Eleven wild type replicates and 11 *uts2r3* mutants were analyzed for the above measures. Outliers from these assessments, defined as >2 SD from the inclusive group mean, were removed following initial review of aggregate data. In addition, due to technical limitations, some specimens did not provide reliable measures of morphology (n = 1), passive mechanics (n = 3), or active contractile function at velocities of 28 Lo/s (n = 8). Active contractile force was not assessed under isometric conditions in three *uts2r3* and one wild type specimen.

Passive mechanical assessments were performed after active mechanics in some specimens, but prior activation did not affect material properties in our comparisons. Before passive testing, a single twitch stimulus (400 mA) was delivered to confirm sample integrity. Data are presented as means ± SD unless otherwise noted. Group-level differences between wild type and *uts2r3* larvae were assessed by ANOVA for active contractile properties, including twitch force, tetanic force, twitch tension, tetanic tension, and velocity-specific deficits. Passive viscoelastic properties were assessed using multivariate ANOVA, with significance measured at each degree of strain. Significance was accepted at *p* < 0.05.

### Micro-computed tomography (µCT) Imaging

Adult zebrafish were euthanized by hypothermic shock and fixed in 4% PFA for 7 days at 4 °C. Fixed specimens were washed twice in 1X PBS to remove residual PFA, then mounted in 0.8% low-melting-point agarose. Specimens were imaged using a ScanCo vivaCT 80 µCT using standard techniques as previously described (20). Three-dimensional reconstructions were generated in 3D Slicer.

